# Inhibition of skeletal muscle Lands cycle ameliorates weakness induced by physical inactivity

**DOI:** 10.1101/2023.07.25.550576

**Authors:** Justin L. Shahtout, Hiroaki Eshima, Patrick J. Ferrara, J. Alan Maschek, James E. Cox, Micah J. Drummond, Katsuhiko Funai

**Affiliations:** Diabetes and Metabolism Research Center, University of Utah, Salt Lake City, UT, USA; Department of Physical Therapy and Athletic Training, University of Utah, Salt Lake City, UT, USA; Molecular Medicine Program, University of Utah, Salt Lake City, UT, USA; Nagasaki International University, Sasebo, Nagasaki, Japan; Metabolomics, Mass Spectrometry, and Proteomics Core, University of Utah, Salt Lake City, UT. USA; Department of Biochemistry, University of Utah, Salt Lake City, UT, USA; Department of Nutrition and Integrative Physiology, University of Utah, Salt Lake City, UT, USA

**Keywords:** skeletal muscle, disuse, atrophy, lipid hydroperoxides, phospholipids

## Abstract

**Background:** Lipid hydroperoxides (LOOH) have been implicated in skeletal muscle atrophy with age and disuse. Lysophosphatidylcholine acyltransferase 3 (LPCAT3), an enzyme of Lands cycle, conjugates a polyunsaturated fatty acyl chain to a lysophospholipid (PUFA-PL) molecule, providing substrates for LOOH propagation. Previous studies suggest that inhibition of Lands cycle is an effective strategy to suppress LOOH. Mice with skeletal muscle-specific tamoxifen-inducible knockout of LPCAT3 (LPCAT3-MKO) were utilized to determine if muscle-specific attenuation of LOOH may alleviate muscle atrophy and weakness with disuse.

**Methods:** LPCAT3-MKO and control mice underwent 7 days of sham or hindlimb unloading (HU model) to study muscle mass and force-generating capacity. LOOH was assessed by quantifying 4-hydroxynonenal (4-HNE)-conjugated peptides. Quantitative PCR and lipid mass spectrometry were used to validate LPCAT3 deletion.

**Results:** 7 days of HU was sufficient to induce muscle atrophy and weakness concomitant to an increase in 4-HNE. Deletion of LPCAT3 reversed HU-induced increase in muscle 4HNE. No difference was found in body mass, body composition, or caloric intake between genotypes. The soleus (SOL) and plantaris (PLANT) muscles of the LPCAT3-MKO mice were partially protected from atrophy compared to controls, concomitant to attenuated decrease in cross-sectional areas in type I and IIa fibers. Strikingly, SOL and extensor digitorum longus (EDL) were robustly protected from HU-induced reduction in force-generating capacity in the LPCAT3-MKO mice compared to controls.

**Conclusion:** Our findings demonstrate that attenuation of muscle LOOH is sufficient to restore skeletal muscle function, in particular a protection from reduction in muscle specific force. Thus, muscle LOOH contributes to atrophy and weakness induced by HU in mice.

## INTRODUCTION

Maintenance of skeletal muscle mass and function is crucial for individuals to maintain their independence and quality of life. Disuse leads to drastic and rapid loss of skeletal muscle mass and strength, of which there is no current pharmacologic therapy [1]. Muscle mass is often very difficult to regain, particularly in our aging population [2]. Increases in reactive oxygen species (ROS) have been implicated as a potential cause for disuse-induced skeletal muscle atrophy [3]. Recently, our laboratory and others have identified the role of a lipid-specific form of ROS known as lipid hydroperoxides (LOOH) in disuse-induced skeletal muscle atrophy [4–7].

LOOH has been previously identified to play a role in the induction of ferroptosis, a non-apoptotic iron-dependent form of regulated cell death [8]. Polyunsaturated fatty acids (PUFAs) provide the essential double bonds in the syntheses of LOOH. Lysophosphatidylcholine acyltransferase 3 (LPCAT3) incorporates PUFAs into lysophospholipids to PUFA containing phospholipids (PUFA-PL). In turn, these PUFAs may be cleaved by phospholipase A_2_ (PLA_2_), reforming a lysophospholipid. This process is referred as the Lands Cycle, responsible for the rearrangements of fatty acyl chains in cellular membrane phospholipids. PUFA-PLs can undergo oxidative attack and lead to the generation of LOOH [9, 10]. Of these, oxidized phosphatidylethanolamine (ox-PE) likely contributes to ferroptosis [11]. Cells contain endogenous LOOH scavenging system by which glutathione peroxidase 4 (GPx4) neutralizes LOOH into nonreactive alcohol [12]. When LOOH propagation exceeds neutralization, LOOH accumulates and begins to form secondary highly reactive molecules that induce carbonyl stress and promote cellular toxicity, including the 9-carbon lipid aldehyde 4-hydroxynonenal (4-HNE).

LOOH has been implicated in a variety of neurological disorders including Alzheimer’s and Parkinson’s disease [13, 14]. Markers of LOOH have been found in various tissues in models of metabolic disease [15–20]. Fewer studies have studied LOOH in skeletal muscle. 4-HNE has been shown to increase in skeletal muscle with long term high fat high sucrose diet in mice [17, 21]. A clinical study also found an increase in 4-HNE in muscle biopsies from individual’s with cancer [22], but others found no increase in 4-HNE with 14 days of lower limb immobilization [23]. Fourteen days of hindlimb immobilization via casting led to increases in 4-HNE protein in plantaris and soleus muscle in mice [24]. Prolonged physical inactivity led to increased skeletal muscle 4-HNE in rats [25]. Multiple studies have confirmed an increase in LOOH in isolated mitochondria and permeabilized fiber bundles in denervated skeletal muscle [26, 27]. Recently, suppression of LOOH via subcutaneous injection of Liproxstatin-1, an inhibitor of LOOH propagation, was found to rescue skeletal muscle atrophy with denervation [4]. Modification of skeletal muscle mitochondrial phospholipids is also sufficient to increase 4-HNE [28, 29].

Our group and others have found that overexpression of GPx4 can neutralize LOOH and/or lipid carbonyls to rescue muscle atrophy and weakness with aging, disuse, and denervation [5, 30, 31]. In these studies, GPx4 overexpression was germline, suggesting that these mice were protected from an increase in LOOH in all their cell-types, not just skeletal muscle [5, 30]. Similarly, treatment of liproxstatin-1, carnosine, or N-acetylcarnosine is effective to suppress LOOH or lipid carbonyls and prevent disuse-induced muscle atrophy [4, 5], but these compounds would be predicted to reduce lipid carbonyl stress throughout the body. Thus, while it may be predicted that muscle LOOH cell-autonomously contribute to atrophy and weakness, there are currently no in vivo data that show this to be the case. The purpose of this study was to test if skeletal muscle-specific modulation of LPCAT3 may rescue LOOH accumulation and prevent disuse-induced muscle atrophy and weakness. Because LPCAT3 deletion occurs only in the skeletal muscle, this mouse model is prevented from LOOH specifically in skeletal muscle.

## METHODS

### Animal Models

To study the effects of LPCAT3 deletion specific to skeletal muscle, the tamoxifen-inducible skeletal muscle-specific knockout (LPCAT3-MKO) mouse was studied, which has been previously utilized by our group [32]. LPCAT3-MKO mice were generated by crossing conditional LPCAT3 knockout (*LPCAT3^fl/fl^*) mice with HSA-MerCreMer mice. Genotypes were determined via PCR. All mice underwent intraperitoneal injection of tamoxifen for 5 consecutive days (7.5 µg/g body mass). After a 2-week washout period from the last day of injection mice then underwent 7 days of HU. All sham and HU mice were between 13-16 weeks of age on the day of tissue harvest. Mice were housed with a 12 hr light/dark cycle in a temperature-controlled room. Mice were fasted for 4 hours prior to anesthetization via intraperitoneal injection of 80 mg/kg ketamine and 10 mg/kg xylazine and tissue harvest. All experimental procedures were approved by the University of Utah Institutional Animal Care and Use Committee.

### Hindlimb Unloading

Mice underwent 7 days of HU with two mice per cage utilizing the same protocol used previously by our group [5, 33]. These mice were checked daily to ensure food and water consumption and to measure body mass. After the 7 days of HU, the mice were fasted for 4 hours then received an intraperitoneal injection of 80 mg/kg ketamine and 10 mg/kg xylazine. After this injection, tissues were harvested. The soleus (SOL), extensor digitorum longus (EDL), gastrocnemius (GAS), tibialis anterior (TA), and plantaris (PLANT) were weighed shortly after dissection.

### Ex Vivo Skeletal Muscle Force Production

Force generating capacity of both the SOL and EDL were measured as previously described [28, 32–34]. Briefly, SOL and EDL were tied with suture at both of their tendons then attached to an anchor and force transducer in a tissue bath (Aurora Scientific, Model 801C) while being submerged in oxygenated Krebs-Henseleit Buffer solution at 37° C. The optimal length of the muscle was then determined via maximum twitch force production. The muscles were then incubated in fresh Krebs-Henseleit Buffer solution and equilibrated for 5 minutes. After this equilibration period, a force-frequency sequence was initiated which stimulated the muscle at increasing frequencies (10, 20, 30, 40, 60, 80, 100, 125, 150, 200 Hz) with a 2-minute rest in between. The rates of contraction and relaxation were quantified as previously described [34]. The Aurora Scientific DMAv5.321 software was utilized for analysis of force production data.

### Mass Spectrometry

All lipidomics analysis was conducted at the Metabolomics Core at the University of Utah. Lipids were extracted from the Tibialis Anterior (TA) muscle with internal standards (Avanti Polar Lipids: 330707) and were analyzed via an Agilent triple-quadrupole time-of-flight mass spectrometer (QTOF-MS/MS). Lipids were isolated using the methyl-*tert*-butyl ether method (MTBE) [32, 34]. Briefly, mouse TA was carefully weighed and homogenized in a methanol/PBS/internal standard solution. MTBE was then added to homogenate and samples were incubated on ice for 60 min and vortexed every 15 min. Samples were centrifuged at 15,000 X *g*, and the organic phase was dried using a Speedvac. The lipids were reconstituted in an 8:2:2 isopropyl alcohol:acetonitrile:ddH_2_O mixture. Phospholipid abundance was normalized to mg of skeletal muscle tissue weight.

### Skeletal Muscle Histology

After careful dissection, the SOL and EDL muscles were embedded in optimal cutting temperature (OCT) gel and sectioned at 10 µm with a cryostat (Microtome Plus). The sections underwent blocking for 1 hr with M.O.M. mouse IgG Blocking Reagent (Vector Laboratories, MKB-2213). Sections were then incubated for 1 hour with concentrated primary antibodies (BA.D5, SC.71, BF.F3 all at 1:100 from DSHB). To detect myosin heavy chain I (MHC I), MHC IIa, and MHC IIb sections were incubated for 1 hr with the following secondary antibodies: Alexa Fluor 647 (1:250; Invitrogen, A21242), Alexa Fluor 488 (1:500; Invitrogen, A21121), and Alexa Fluor 555 (1:500; Invitrogen, A21426). Negative stained fibers were considered to be IIx. After staining, slides were mounted with mounting medium (Vector Laboratories, H-1000). Slides were imaged with an automated wide-field light microscope (Nikon Corp.) utilizing a 10x objective lens. The cross-sectional area of these fibers was then quantified utilizing ImageJ software.

### Western blotting

Whole muscle lysate was utilized for western blotting. Frozen GAS muscle was homogenized in a glass homogenization tube using a mechanical pestle grinder with homogenization buffer (50mM Tris pH 7.6 5 mM EDTA 150mM NaCl 0.1% SDS 0.1% sodium deoxycholate 1% triton X-100 protease inhibitor cocktail). After homogenization, samples were centrifuged for 15 min at 12,000 X *g*. Protein concentration of supernatant was then determined using a BCA protein Assay Kit (Thermo Scientific). Equal protein was then mixed with Laemmeli sample buffer and loaded onto 4-20% gradient gels (Bio-Rad). Transfer of proteins occurred on a nitrocellulose membrane which was then Ponceau S. stained and imaged for equal protein loading and transfer quality. Membranes were then blocked for 1 hr. at room temperature with 5% bovine serum albumin in Tris-buffered saline with 0.1% Tween 20 (TBST). The membranes were then incubated with primary antibody for GAPDH (Cell Signaling Technology, 14C10) and 4-HNE (abcam, ab48506). Membranes were imaged utilizing Western Lightning Plus-ECL (PerkinElmer) and a FluorChem E Imager (Protein Simple). Image Lab software (BioRad) was used for analysis of western blot images.

### Quantitative reverse transcription PCR

TA muscle samples were homogenized in TRIzol (Life Technologies) for total RNA extraction. One microgram of RNA was reverse-transcribed using an IScript cDNA synthesis kit (Bio-Rad). Reverse transcription PCR (RT-PCR) was performed with the Viia 7 Real-Time PCR System (Life Technologies) using a SYBR Green reagent (Life Technologies). Data was then normalized to the ribosomal protein L32 gene expression levels and then normalized to the mean of the control group. The following LPCAT3 and L32 primers were used: LPCAT3-F (GGCCTCTCAATTGCTTATTTCA) LPCAT3-R (AGCACGACACATAGCAAGGA) L32-F (TTCCTGGTCCACAATGTCAA) L32-R (GGCTTTTCGGTTCTTAGAGGA).

### Statistical Analyses

Values are presented as means ± SEM. Analyses were performed using GraphPad Prism 9.1.1 software. Statistical comparisons were made using an unpaired two-tailed Student’s t test for two groups. For more than two groups, a two-way analysis of variance (ANOVA) was performed followed by the appropriate multiple-comparison test. For all tests, P value less than 0.05 was considered statistically significant.

## RESULTS

### 7 days of hindlimb unloading increases skeletal muscle lipid hydroperoxides

Hindlimb unloading (HU) decreased body mass and fat mass with no significant detectable change in lean mass (Figure 1A-D). Nonetheless, HU led to a ~20% decrease in soleus (SOL) mass with a drastic decrease in muscle force production (Figure 1E-F). HU did not affect extensor digitorum longus (EDL) mass but robustly decreased force-generating capacity (Figure 1G-H). As previously described [5, 33], HU led to a robust accumulation of 4-hydroxnonenal (4-HNE) in skeletal muscle (Figure I-J).

**Figure 1.**
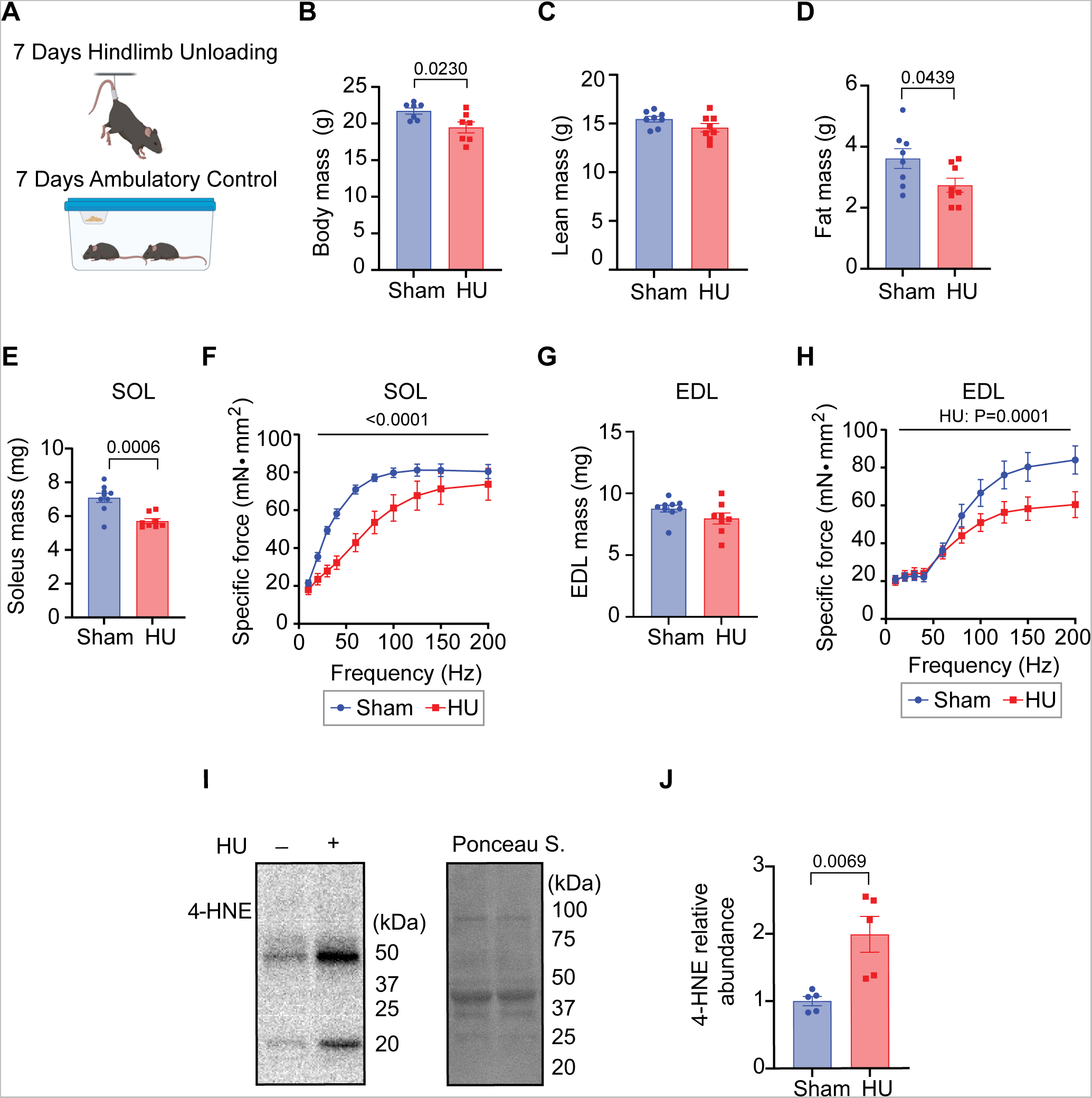
Hindlimb unloading (HU) increases lipid hydroperoxides (LOOH) in mouse skeletal muscle. A) Schematic of HU and ambulatory control (sham) housing conditions. B-D) Body mass (B), lean mass (C), and fat mass (D) of sham and HU mice (*n* = 7/group). E-F) Soleus (SOL) mass (*n* = 8-9/group) (E) and force frequency curve (sham *n* = 6, HU *n* = 5) (F). G-H) Extensor digitorum longus (EDL) mass (*n =* 8-9/group) (G) and force frequency curve (sham *n* = 6, HU *n* = 5) (H). I-J) 4-hydroxynonenal (4-HNE) representative western blot image from skeletal muscle homogenate with Ponceau S. imaged as a protein loading control (I) and quantification (*n* = 5/group) (J). For (B-E,G,J) unpaired two-tailed t-tests were performed. For (F,H) two-way ANOVA were performed with Sidak’s post-hoc test. All data are mean ± SEM.

### LPCAT3 deletion prevents muscle lipid hydroperoxide accumulation induced by HU

After tamoxifen injections and a 2-wk washout, LPCAT3-MKO and control mice were subject to HU or sham treatment for 7 days. All measurements were taken on terminal day (Figure 2A). As expected, *LPCAT3* transcripts were significantly reduced in skeletal muscle of LPCAT3-MKO mice compared to controls regardless of the HU intervention. HU also reduced skeletal muscle *LPCAT3* transcript (Figure 2B). Consistent with the notion that PUFA incorporation is essential for LOOH propagation, HU-induced increase in skeletal muscle 4-HNE was completely reversed with muscle-specific LPCAT3 deletion (Figure 2C-D).

**Figure 2.**
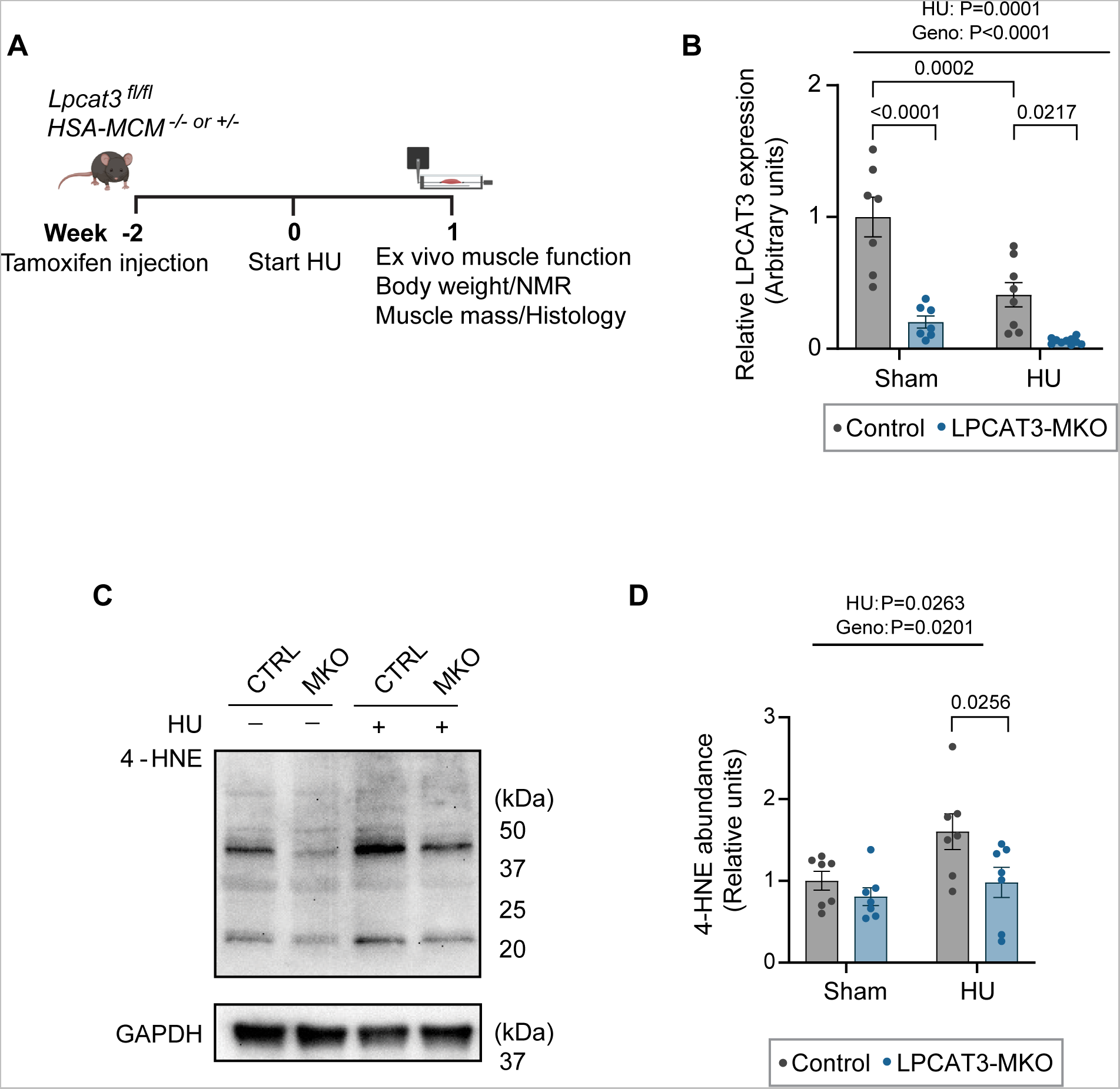
Skeletal muscle specific knockout of lysophosphatidylcholine acyltransferase 3 (LPCAT3-MKO) prevents skeletal muscle lipid hydroperoxide (LOOH) accumulation with hindlimb unloading (HU). A) Schematic of experimental timeline. B) Skeletal muscle LPCAT3 mRNA abundance in control and LPCAT3-MKO mice in both the sham and HU groups (*n* = 7-10/group). C-D) 4-hydroxynonenal (4-HNE) and GAPDH representative western blot image from skeletal muscle homogenate (C) and quantification (D) (*n* = 7/group). Two-way ANOVA was performed with Sidak’s post-hoc test. All data are mean ± SEM.

### Skeletal muscle LPCAT3 knockout only alters PUFA-PL species in sham groups

Sham LPCAT3-MKO mice displayed lower values of the most abundant PUFA-PC species compared to control mice, specifically 16:0,20:4 PUFA-PC (Figure 3A). Similar to a previous finding [35], a small but not robust difference was observed in LysoPC levels in sham mice (Figure 3B). Consistent with LPCAT3’s activity towards lyso-PE in addition to lyso-PC, multiple PUFA-PE species were decreased in the LPCAT3-MKO sham mice (16:0,20:4, 16:0,22:6, 18:0,20:4, and 18:2,22:4) (Figure 3C). Strikingly, no significant differences were observed in PC, LysoPC, and LysoPE between genotypes after HU (Figures 3E,F,H). Noticeable increases in few PUFA-PE species were observed in the LPCAT3-MKO mice with HU (18:1,22:6 and 18:2,22:6) (Figure 3G). We take these findings to mean that HU decelerates skeletal muscle Lands cycle such that changes in flux through Lands cycle is not reflected in a static measurement of phospholipids. Reduced muscle LPCAT3 transcript with HU is consistent with this idea (Figure 2B).

**Figure 3.**
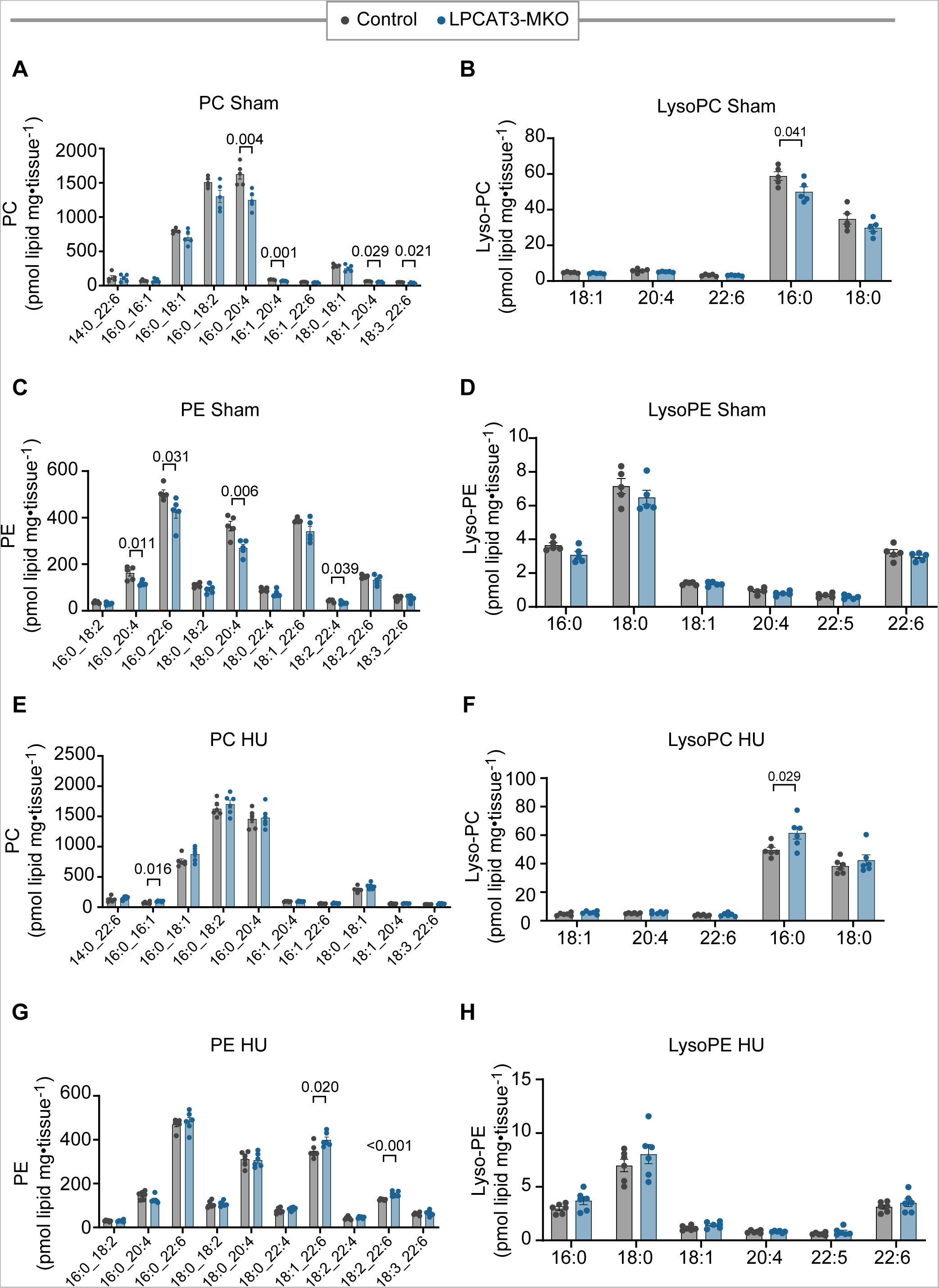
Skeletal muscle specific knockout of lysophosphatidylcholine acyltransferase 3 (LPCAT3-MKO) decreases PUFA-PC only in the sham group. A-D) Skeletal muscle phosphatidylcholine (PC) (A), lysophosphatidylcholine (LysoPC) (B), phosphatidylethanolamine (PE) (C), lysophosphatidylethanolamine (LysoPE) (D) abundance in sham control and LPCAT3-MKO mice (*n* = 5/group). E-H) Skeletal muscle phosphatidylcholine (PC) (E), lysophosphatidylcholine (LysoPC) (F), phosphatidylethanolamine (PE) (G), lysophosphatidylethanolamine (LysoPE) (H) abundance of hindlimb unloading (HU) control and LPCAT3-MKO mice (*n* = 6/group). Unpaired two-tailed t-tests were performed. All data are mean ± SEM.

### Muscle-specific LPCAT3 deletion does not alter body composition with or without HU

Seven days of HU was sufficient to decrease body and lean masses regardless of genotypes (Figure 4A,B). Hindlimb unloading had increased fat mass regardless of genotypes, likely reflecting reduced energy expenditure (Figure 4C). No differences in food intake was found between genotypes with hindlimb unloading (Figure 4D).

**Figure 4.**
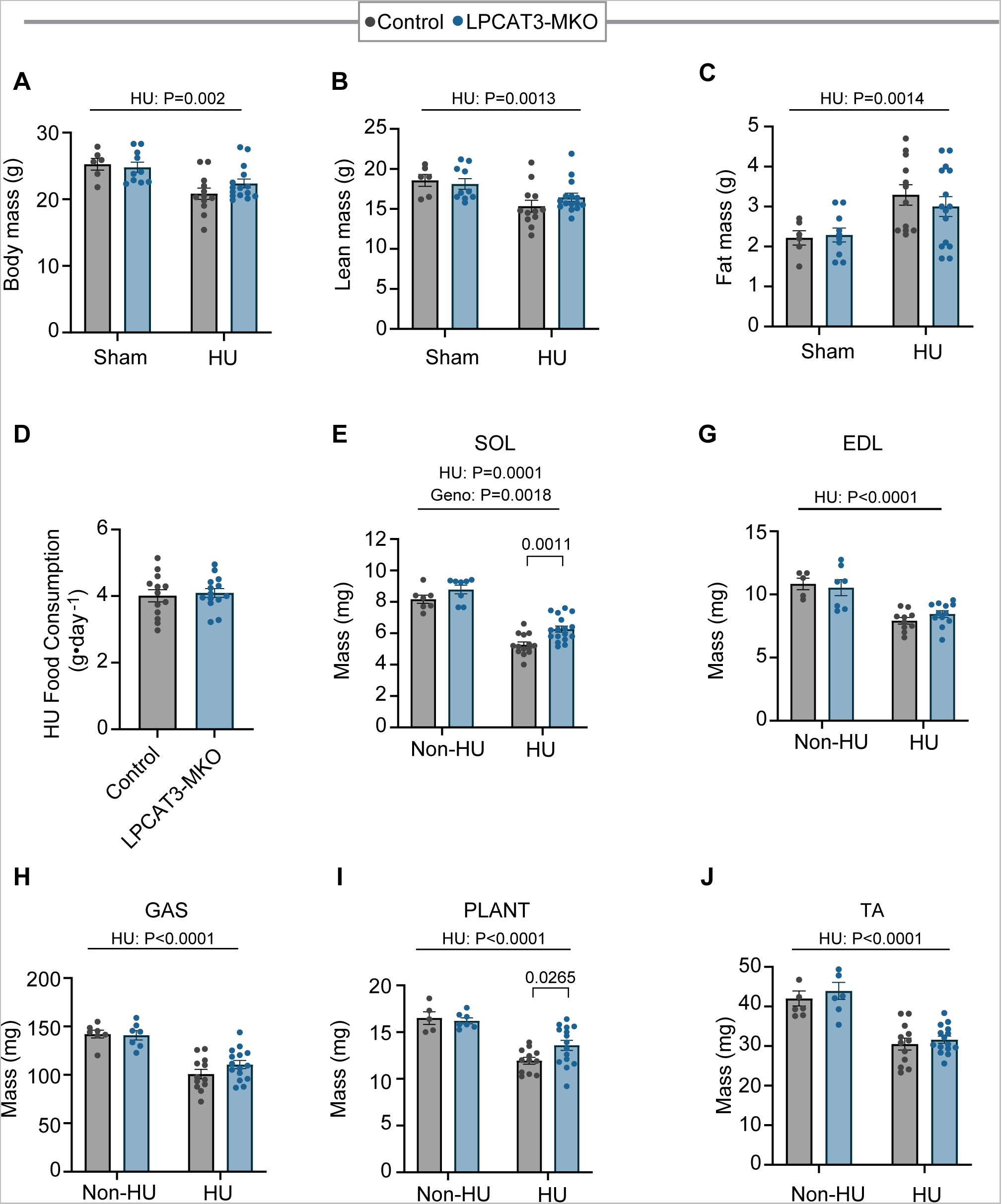
Skeletal muscle specific knockout of lysophosphatidylcholine acyltransferase 3 (LPCAT3-MKO) does not alter body composition but rescues skeletal muscle atrophy with hindlimb unloading (HU). A-C) Bodyweight (A), lean mass (B), and fat mass (C) from control and LPCAT3-MKO mice in sham and HU groups (*n* = 6-15/per group). D) Average food consumption of control and LPCAT3-MKO mice across the 7 days of HU (*n* = 13-14/group). E-J) Mass of several hindlimb muscles from control and LPCAT3-MKO mice in sham and HU groups (*n* = 7-17/group). Two-way ANOVA was performed with Sidak’s post-hoc test. All data are mean ± SEM. For (A-C) and (E-J) two-way ANOVA was performed with Sidak’s post-hoc test. Unpaired t-test was performed in (D). All data are mean ± SEM.

### LPCAT3 deletion partially protects from skeletal muscle atrophy induced by HU

HU led to drastic decreases in mass of the SOL, gastrocnemius (GAS), EDL, plantaris (PLANT), and tibialis anterior (TA) muscles (Figure 4E-J). This effect was most pronounced in the SOL muscle, where the LPCAT3-MKO mouse displayed higher mass with and without hindlimb unloading (HU) (Figure 4E). LPCAT3-MKO mice experience ~13% less SOL atrophy than control mice with HU. The PLANT muscle of the KO mice also experienced ~43% less atrophy compared to genotype controls (Figure 4I).

### LPCAT3 knockout protects against HU-induced decreases in myofiber cross-sectional areas in type I and IIa fibers

Consistent with previous work, SOL muscle was comprised of type I and type IIa muscle fibers [5, 33]. 7 days of HU led to an approximately 40% decrease in type I and type IIa fiber CSA in the SOL muscle. This was decreased to only ~15% in the LPCAT3-KO mice in both of these fiber types (Figure 5A-C). EDL was comprised of type IIa, IIb, and IIx fibers with a negligible amount of type I fibers. HU led to a significant decrease in CSA of type IIb fibers of the EDL, yet no differences were found between genotype (Figure 5D-F). 7 days of hindlimb unloading was not sufficient to cause changes in muscle fiber type composition of SOL or EDL. This is consistent with previous use of the 7-day HU model by our group [33].

**Figure 5.**
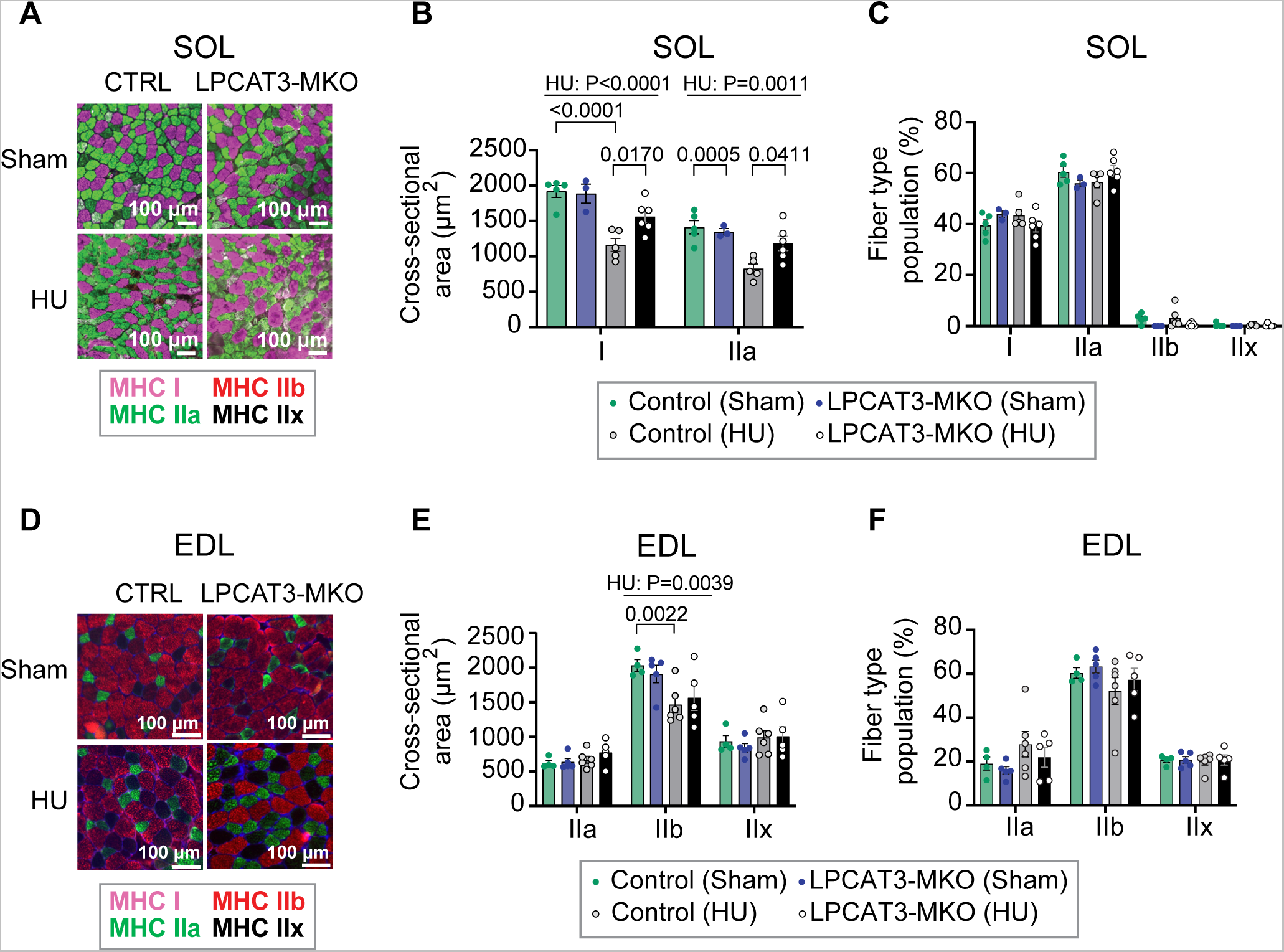
Skeletal muscle specific knockout of lysophosphatidylcholine acyltransferase 3 (LPCAT3-MKO) rescues myofiber cross-sectional area in soleus (SOL) muscle with hindlimb unloading (HU). A) Representative myosin heavy chain (MHC) isoform immunofluorescence images of SOL muscles from control and LPCAT3-MKO mice in sham and HU groups. Scale bar is 100µm. B) Muscle fiber cross-sectional area by fiber type of SOL muscles from control and LPCAT3-MKO mice in sham and HU groups (*n =* 3-6/group). C) Muscle fiber type composition of SOL muscles from control and LPCAT3-MKO mice in sham and HU groups (*n =* 3-6/group). D) Representative MHC isoform immunofluorescence images of extensor digitorum longus (EDL) muscles from control and LPCAT3-MKO mice in sham and HU groups. Scale bar is 100µm. E) Muscle fiber cross-sectional area by fiber type of EDL muscles from control and LPCAT3-MKO mice in sham and HU groups (*n =* 4-6/group). C) Muscle fiber type composition of EDL muscles from control and LPCAT3-MKO mice in sham and HU groups (*n =* 4-6/group). Two-way ANOVA was performed with Sidak’s post-hoc test. All data are mean ± SEM.

### LPCAT3 knockout robustly protects musckes from the loss of force-generating capacity induced by HU

Both SOL and EDL muscles underwent decreases in twitch forces with HU (Figure 6A-D). Strikingly, both of these muscles experienced a significant rescue in twitch force production in the KO mice, nearly back to sham levels. Similarly, both SOL and EDL muscles were protected from HU-induced reduction in tetanic force production. In SOL, force-generating capacity was completely rescued in LPCAT3-MKO mice back to the sham level (Figure 6E,F). In EDL, LPCAT3-MKO mice were partly rescued from disuse-induced muscle weakness (Figure 6G,H), similar to data on muscle atrophy (Figure 4E). To further examine the muscle function in these mice, average rate of contraction and relaxation as well as the stimulation frequency needed to illicit 50% maximum force production (IC_50_) values were calculated. The SOL average rate of contraction was significantly rescued in the KO mouse, with the average rate of relaxation trending (Figure 7A,B). As expected, HU drastically decreased the SOL muscle’s sensitivity to stimulation as seen by an increase in IC_50_, but no differences in genotype (Figure 7C). HU decreased the average rate of contraction and relaxation in the EDL, but no differences between genotype were found (Figure 7D,E). Neither HU nor genotype influenced the EDL’s sensitivity to stimulation (Figure 7F).

**Figure 6.**
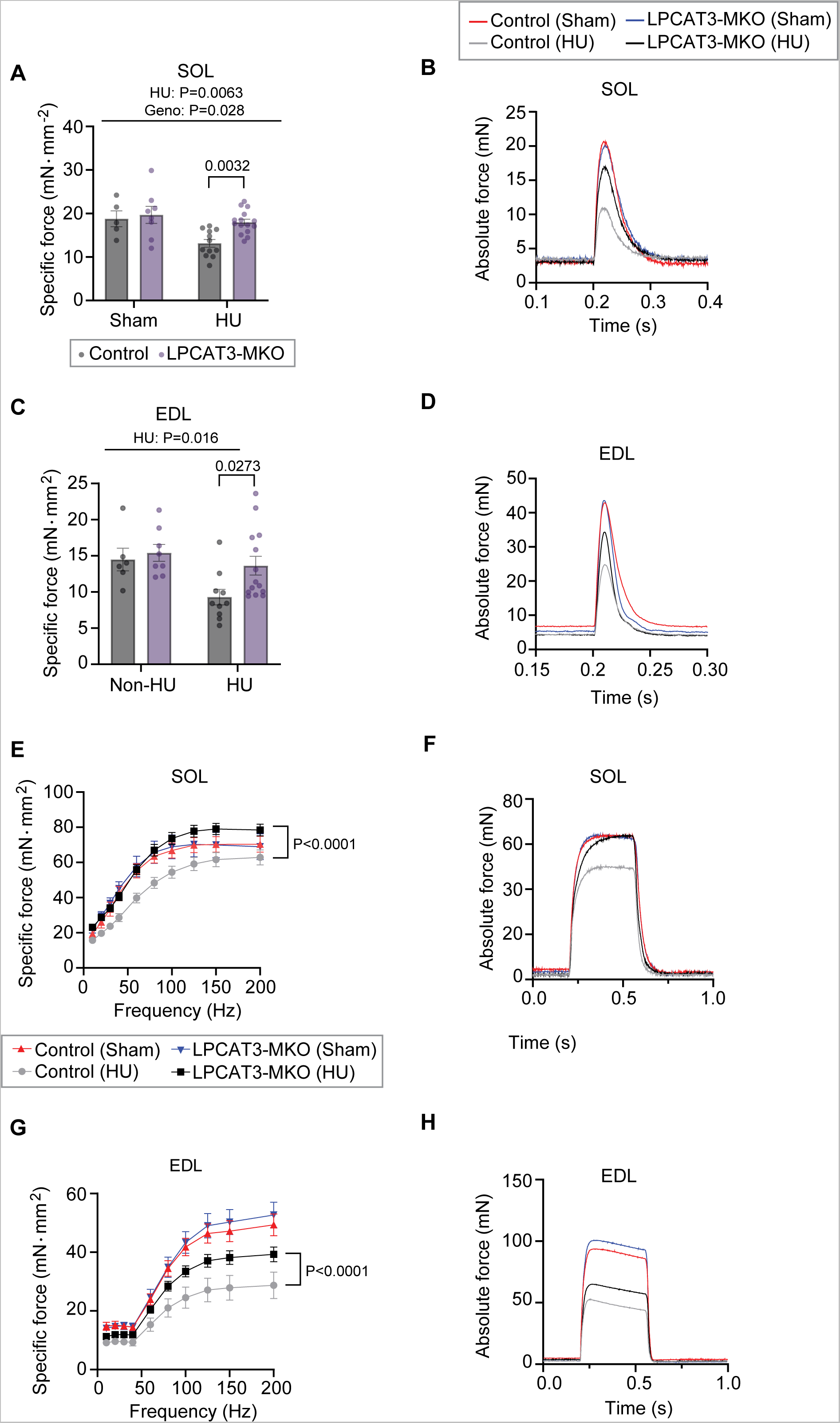
Skeletal muscle specific knockout of lysophosphatidylcholine acyltransferase 3 (LPCAT3-MKO) rescues skeletal muscle weakness with hindlimb unloading (HU). A) Soleus (SOL) twitch force production from control and LPCAT3-MKO mice in sham and HU groups (*n* = 5-15/group). B) SOL representative twitch force tracings from control and LPCAT3-MKO mice in sham and HU groups. C) Extensor digitorum longus (EDL) twitch force production from control and LPCAT3-MKO mice in both sham and HU groups (*n* = 6-16/group). D) EDL representative twitch force tracings from control and LPCAT3-MKO mice in sham and HU groups. E) Force frequency curves from SOL muscles from control and LPCAT3-MKO mice in sham and HU groups (control sham *n* = 6, LPCAT3-MKO sham *n* = 8, control HU *n* = 12, LPCAT3-MKO HU *n* = 12). F) Representative tetanic tracings from SOL muscles from control and LPCAT3-MKO mice in both sham and HU groups. G) Force frequency curves from EDL muscles from control and LPCAT3-MKO mice in sham and HU groups (control sham *n* = 8, LPCAT3-MKO sham *n* = 9, control HU *n* = 11, LPCAT3-MKO HU *n* = 15). H) Representative tetanic tracings from EDL muscles from control and LPCAT3-MKO mice in both sham and HU groups. Two-way ANOVA was performed with Sidak’s post-hoc test. All data are mean ± SEM.

**Figure 7.**
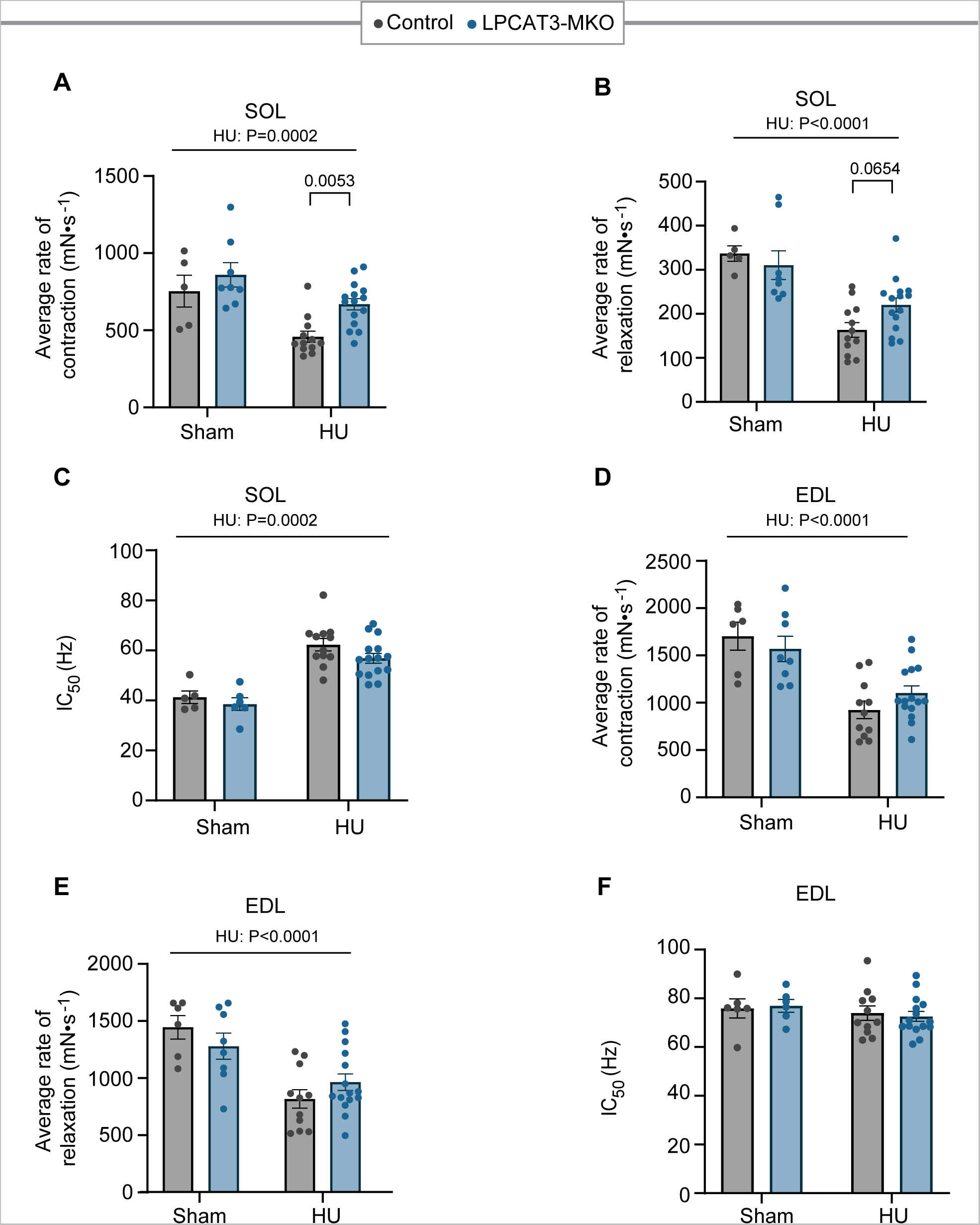
Skeletal muscle specific knockout of lysophosphatidylcholine acyltransferase 3 (LPCAT3-MKO) rescues soleus (SOL) muscle contractile dynamics with hindlimb unloading (HU). A) Average rate of contraction of SOL muscles from control and LPCAT3-MKO mice in sham and HU groups (*n* = 5-15/group). B) Average rate of relaxation of SOL muscles from control and LPCAT3-MKO mice in sham and HU groups (*n* = 5-15/group). C) IC_50_ values of SOL muscles from control and LPCAT3-MKO mice in sham and HU groups (*n* = 5-15/group). D) Average rate of contraction of extensor digitorum longus (EDL) muscles from control and LPCAT3-MKO mice in sham and HU groups (*n =* 6-15/group). E) Average rate of relaxation in EDL muscles from control and LPCAT3-MKO mice in sham and HU groups (*n =* 6-15/group). F) IC_50_ values of EDL muscles from control and LPCAT3-MKO mice in sham and HU groups (*n =* 6-15/group). Two-way ANOVA was performed with Sidak’s post-hoc test. All data are mean ± SEM.

## DISCUSSION

Disuse leads to drastic skeletal muscle atrophy and weakness [1]. ROS has been implicated as a mediator in this process, yet the exact subspecies of ROS responsible for these mechanisms are not fully understood [3]. Previous work by our group and others suggests lipid hydroperoxides act to promote disuse-induced skeletal muscle atrophy [4–6, 30, 31]. Mice displaying whole-body overexpression of GPx4 display protection from disuse- and denervation-induced atrophy, providing strong evidence that an increase in LOOH scavenging promotes beneficial effects on skeletal muscle [5, 30, 31]. However, to our knowledge, there has not been studies where LOOH was suppressed specifically in skeletal muscle to show cell-autonomous protection from disuse-induced atrophy *in-vivo*. To address this question, we interrogated the Lands cycle specifically in skeletal muscle by performing skeletal muscle-specific knockout of LPCAT3, which would be predicted to inhibit the supply chain for fatty acid double bonds that are essential for LOOH propagation. Indeed, LPCAT3-MKO mice were protected from disuse-induced increase in skeletal muscle 4-HNE. Importantly, these mice were partially protected from disuse-induced muscle atrophy and weakness.

Lands cycle, in particular LPCAT3-dependent acylation, plays an important role in incorporation of PUFA in the membrane phospholipids. LPCAT3 displays high affinity towards 18:2 and 20:4 acyl-chains to be incorporated into lysophospholipids. The high affinity of LPCAT3 towards PUFA incorporation makes this enzyme an essential enzyme also for lipid peroxidation. Indeed, muscle-specific deletion of LPCAT3 appeared to effectively suppressed accumulation of LOOH, demonstrated by prevention of 4-HNE accumulation with HU. Some studies indicate that oxPE is the important LOOH molecule that signal to ferroptosis [11, 36], though it is uncertain whether mechanisms for ferroptosis overlaps with mechanisms that promote muscle atrophy and weakness with aging or disuse. Nevertheless, multiple downstream fates of oxidized phospholipids have been implicated in cellular damage. Oxidized acyl chains may degrade into highly reactive lipid carbonyl end products, such as 4-HNE. These lipid carbonyls covalently bind to proteins, altering their function, and even promoting their degradation [37]. Our previously study shows neutralization of lipid carbonyl stress with N-acetylcarnosine treatment or GPx4 overexpression effectively suppresses 4-HNE accumulation, and protects from muscle atrophy and weakness with age and disuse [5]. Very similar and parallel results were shown by the Van Remmen group, demonstrating that GPx4 overexpression, lipoxstatin-1 treatment, and inhibition of cPLA2 are sufficient to prevent against denervation- or aging-induced muscle LOOH accumulation and atrophy [4, 6, 30, 31, 38]. Instead of lipid carbonyls, they implicate oxylipins another class of lipids downstream of LOOH to decline in muscle function. Thus, the exact mechanism downstream of LOOH that promote disuse-induced muscle atrophy remains uncertain.

Low LPCAT3 levels led to decrease in PUFA-PC species only in the sham group with no changes between genotype with HU. While unexpected, these data resemble previous work in the liver and muscle, which demonstrated that high-fat diet may accelerate Lands cycle and the turnover of PUFA-PL compared to standard chow [32, 35]. Similarly, changes in PUFA-PL and lyso-PL levels with LPCAT3 deletion was more modest in our standard-chow fed sham group. The differences between in the lipidome control and LPCAT3-MKO mice were further diminished with hindlimb unloading. Together with findings that HU itself reduced muscle LPCAT3 level, we interpret these data to mean that HU decelerates Lands cycle such that deletion of LPCAT3 is not reflected in static measurements of PC or lyso-PC.

It is difficult to conclude whether there were fiber-type specific effects on rescuing muscle mass or force-generating capacity. LPCAT3-MKO mice were partially protected from skeletal muscle atrophy in the SOL and PLANT muscles, but not in EDL, GAS, or TA muscles. SOL is predominantly slow and PLANT and EDL are predominantly fast, with GAS and TA having mixed constituents. In SOL muscles, fiber CSA were partially restored in both type I and IIa fibers (no IIx fibers detected), while in EDL muscles, CSA were not different for IIa fibers in addition to IIb and IIx fibers. Nevertheless, HU induced reduction in CSA in both I and IIa fibers in SOL, but only in IIb fibers in EDL. Meanwhile, ex vivo force production were rescued in both SOL and EDL muscles, but to a greater degree in SOL than EDL. Because type I/IIa fibers are more oxidative than type IIb/IIx fibers, it is reasonable to suspect that neutralization of LOOH have greater protective effects in type I/IIa fibers for both mass and force.

Our data demonstrate that neutralization of muscle LOOH more robustly restores muscle force than mass, similar to our previously work [5]. These findings suggest that LOOH, and potential the lipid carbonyls, likely disproportionately affect enzymes involved in regulating muscle contraction, excitation-contraction coupling, or energy metabolism rather than enzymes involved in protein synthesis or degradation. It is interesting to note that a recent study indicates that defects in muscle energetics and force-generating capacity may precede muscle atrophy in cachexia, consistent with the idea that a separate set of mechanisms contributes to reductions in muscle mass and force [39].

In conclusion, the current study shows that skeletal muscle-specific suppression of LOOH can protect muscles from disuse-induced atrophy and weakness. Similar to studies using global suppression of LOOH, the effect was more robust in preserving the force-generating capacity of muscles than in preserving mass. Other studies have shown therapeutic potential for targeting this pathway to treat muscle atrophy [4–6, 30, 31, 38]. There are further questions regarding the role of LOOH in muscle atrophy and weakness that remain unanswered. In particular, what are the downstream mechanisms of LOOH? What skeletal muscle proteins are most susceptible to carbonylation and how does this influence skeletal muscle function? Future studies should address how LOOH affects the skeletal muscle proteome to influence their activities.

## Nonstandard Abbreviations

EDL: Extensor digitorum longus
GAS: Gastrocnemius
GPx4: Glutathione peroxidase 4
HU: Hindlimb unloading
SOL: Soleus
LPCAT3: Lysophosphatidylcholine acyltransferase 3
LysoPC: Lysophosphatidylcholine
LysoPE: Lysophosphatidylethanolamine
MKO: Muscle-specific knockout
PLA_2_: Phospholipase A2
PUFA-PL: Polyunsaturated acyl chain phospholipid
PC: Phosphatidylcholine
Phosphatidylethanolamine
LOOH: Lipid hydroperoxides
MS/MS: Tandem mass spectrometry
ox-PE: oxidized phosphatidylethanolamine
PLANT: Plantaris
QTOF: Quadrupole time-of-flight
TA: Tibialis anterior
4-HNE: 4-hydroxynonenal

## Acknowledgements

This research is supported by NIH grants DK107397, DK127979, GM144613 (to KF), AG074535 (to KF and MJD), and AG076075 (to MJD), Larry H. and Gail Miller Family Foundation (to PJF), Uehara Memorial Foundation (to HE), University of Utah Center on Aging Pilot Grant (to KF), S10 OD016232, S10 OD021505, and U54 DK110858 to University of Utah Metabolomics Core Facility. We would like to thank Dr. Peter Tontonoz from the University of California, Los Angeles for graciously sharing the LPCAT3 conditional knockout mouse. We would also like to thank Nikita Abraham and Diana Lim from the University of Utah Molecular Medicine Program for assistance with figures.

## Author Contributions

JLS, KF, HE, PJF, MJD contributed to the study design. JLS and KF wrote the manuscript. JLS and HE performed ex vivo contraction. JLS performed qPCR, western blotting, immunohistochemistry, and lipid extraction experiments. KF, PJF, and JLS developed and maintained the mouse colony. JAM and JEC performed mass spectrometry analyses.

## References

1. Bodine, S.C., Disuse-induced muscle wasting. Int J Biochem Cell Biol, 2013. 45(10): p. 2200–8.

2. Wall, B.T., M.L. Dirks, and L.J. van Loon, Skeletal muscle atrophy during short-term disuse: implications for age-related sarcopenia. Ageing Res Rev, 2013. 12(4): p. 898–906.

3. Powers, S.K., A.J. Smuder, and D.S. Criswell, Mechanistic links between oxidative stress and disuse muscle atrophy. Antioxid Redox Signal, 2011. 15(9): p. 2519–28.

4. Brown, J.L., et al., Lipid hydroperoxides and oxylipins are mediators of denervation induced muscle atrophy. Redox Biology, 2022. 57.

5. Eshima, H., et al., Lipid hydroperoxides promote sarcopenia through carbonyl stress. Elife, 2023. 12.

6. Pharaoh, G., et al., Targeting cPLA2 derived lipid hydroperoxides as a potential intervention for sarcopenia. Scientific Reports, 2020. 10(1).

7. Miranda, E.R., J.L. Shahtout, and K. Funai, Chicken or egg? Mitochondrial phospholipids and oxidative stress in disuse-induced skeletal muscle atrophy. Antioxid. Redox Signal., 2022.

8. Dixon, S.J., et al., Ferroptosis: an iron-dependent form of nonapoptotic cell death. Cell, 2012. 149(5): p. 1060–72.

9. Lands, W.E., Metabolism of glycerolipides; a comparison of lecithin and triglyceride synthesis. The Journal of biological chemistry, 1958. 231(2): p. 883–888.

10. Tyurina, Y.Y., et al., “Only a Life Lived for Others Is Worth Living”: Redox Signaling by Oxygenated Phospholipids in Cell Fate Decisions. Antioxid Redox Signal, 2018. 29(13): p. 1333–1358.

11. Kagan, V.E., et al., Oxidized arachidonic and adrenic PEs navigate cells to ferroptosis. Nature Chemical Biology, 2017. 13(1): p. 81–90.

12. Yang, W.S., et al., Regulation of ferroptotic cancer cell death by GPX4. Cell, 2014. 156(1-2): p. 317–331.

13. Chen, L., et al., Ablation of the ferroptosis inhibitor glutathione peroxidase 4 in neurons results in rapid motor neuron degeneration and paralysis. Journal of Biological Chemistry, 2015. 290(47): p. 28097–28106.

14. Do Van, B., et al., Ferroptosis, a newly characterized form of cell death in Parkinson’s disease that is regulated by PKC. Neurobiol Dis, 2016. 94: p. 169–78.

15. Pillon, N.J., et al., Nonenal (4-HNE) Induces Insulin Resistance in Skeletal Muscle through Both Carbonyl and Oxidative Stress. 2012. 153(May): p. 2099–2111.

16. Soulage, C.O., et al., Skeletal muscle insulin resistance is induced by 4-hydroxy-2-hexenal, a by-product of n −3 fatty acid peroxidation. 2018: p. 688–699.

17. Anderson, E.J., et al., A carnosine analog mitigates metabolic disorders of obesity by reducing carbonyl stress. Journal of Clinical Investigation, 2018. 128(12): p. 5280–5293.

18. Katunga, L.A., et al., Obesity in a model of gpx4 haploinsufficiency uncovers a causal role for lipid-derived aldehydes in human metabolic disease and cardiomyopathy. Mol Metab, 2015. 4(6): p. 493–506.

19. Samjoo, I.A., et al., The effect of endurance exercise on both skeletal muscle and systemic oxidative stress in previously sedentary obese men. Nutr Diabetes, 2013. 3: p. e88.

20. Aldini, G., et al., The carbonyl scavenger carnosine ameliorates dyslipidaemia and renal function in Zucker obese rats. J Cell Mol Med, 2011. 15(6): p. 1339–54.

21. Lee, H., et al., Mitochondrial dysfunction in skeletal muscle contributes to the development of acute insulin resistance in mice. J Cachexia Sarcopenia Muscle, 2021. 12(6): p. 1925–1939.

22. Brzeszczyńska, J., et al., Loss of oxidative defense and potential blockade of satellite cell maturation in the skeletal muscle of patients with cancer but not in the healthy elderly. Aging (Albany NY), 2016. 8(8): p. 1690–1702.

23. Miotto, P.M., et al., Supplementation with dietary ω-3 mitigates immobilization-induced reductions in skeletal muscle mitochondrial respiration in young women. FASEB Journal, 2019. 33(7): p. 8232–8240.

24. Min, K., et al., Mitochondrial-targeted antioxidants protect skeletal muscle against immobilization-induced muscle atrophy. 2022. 32611: p. 1459–1466.

25. Yoshihara, T., et al., Long-term physical inactivity exacerbates hindlimb unloading-induced muscle atrophy in young rat soleus muscle. J Appl Physiol (1985), 2021. 130(4): p. 1214–1225.

26. Muller, F.L., et al., Denervation-induced skeletal muscle atrophy is associated with increased mitochondrial ROS production. Am J Physiol Regul Integr Comp Physiol, 2007. 293(3): p. R1159–68.

27. Sataranatarajan, K., et al., Molecular changes in transcription and metabolic pathways underlying muscle atrophy in the CuZnSOD null mouse model of sarcopenia. Geroscience, 2020. 42(4): p. 1101–1118.

28. Heden, T., et al., Mitochondrial PE potentiates respiratory enzymes to amplify skeletal muscle aerobic capacity. Science Advances, 2019. eaax8352.

29. Johnson, J.M., et al., Targeted overexpression of catalase to mitochondria does not prevent cardioskeletal myopathy in Barth syndrome. J Mol Cell Cardiol, 2018. 121: p. 94–102.

30. Bhattacharya, A., et al., Denervation induces cytosolic phospholipase A2-mediated fatty acid hydroperoxide generation by muscle mitochondria. J Biol Chem, 2009. 284(1): p. 46–55.

31. Czyżowska, A., et al., Elevated phospholipid hydroperoxide glutathione peroxidase (GPX4) expression modulates oxylipin formation and inhibits age-related skeletal muscle atrophy and weakness. Redox Biology, 2023. 64.

32. Ferrara, P.J., et al., Lysophospholipid acylation modulates plasma membrane lipid organization and insulin sensitivity in skeletal muscle. Journal of Clinical Investigation, 2021. 131(8).

33. Eshima, H., et al., Neutralizing mitochondrial ROS does not rescue muscle atrophy induced by hindlimb unloading in female mice. J Appl Physiol (1985), 2020. 129(1): p. 124–132.

34. Ferrara, P.J., et al., Low lysophosphatidylcholine induces skeletal muscle myopathy that is aggravated by high-fat diet feeding. FASEB J, 2021. 35(10): p. e21867.

35. Rong, X., et al., Lpcat3-dependent production of arachidonoyl phospholipids is a key determinant of triglyceride secretion. 2015: p. 1–23.

36. Bayir, H., et al., Ferroptotic mechanisms and therapeutic targeting of iron metabolism and lipid peroxidation in the kidney. Nat Rev Nephrol, 2023. 19(5): p. 315–336.

37. Castro, J.P., et al., 4-Hydroxynonenal (HNE) modified proteins in metabolic diseases. Free Radic Biol Med, 2017. 111: p. 309–315.

38. Bhattacharya, A., et al., Genetic ablation of 12/15-lipoxygenase but not 5-lipoxygenase protects against denervation-induced muscle atrophy. Free Radic Biol Med, 2014. 67: p. 30–40.

39. Delfinis, L.J., et al., Muscle weakness precedes atrophy during cancer cachexia and is linked to muscle-specific mitochondrial stress. JCI Insight, 2022. 7(24).

